# Performance Monitoring for Sensorimotor Confidence: A Visuomotor Tracking Study

**DOI:** 10.1101/861302

**Authors:** Shannon M. Locke, Pascal Mamassian, Michael S. Landy

**Affiliations:** Laboratoire des Systèmes Perceptifs, Département d’Études Cognitives, École Normale Supérieure, PSL University, CNRS, 75005 Paris, France; Department of Psychology, New York University, New York, NY, United States; Center for Neural Science, New York University, New York, NY, United States

**Keywords:** sensorimotor, confidence, metacognition, perception, action, tracking

## Abstract

To best interact with the external world, humans are often required to consider the quality of their actions. Sometimes the environment furnishes rewards or punishments to signal action efficacy. However, when such feedback is absent or only partial, we must rely on internally generated signals to evaluate our performance (i.e., metacognition). Yet, very little is known about how humans form such judgements of sensorimotor confidence. Do they monitor their performance? Or do they rely on cues to sensorimotor uncertainty to infer how likely it is they performed well? We investigated motor metacognition in two visuomotor tracking experiments, where participants followed an unpredictably moving dot cloud with a mouse cursor as it followed a random trajectory. Their goal was to infer the underlying target generating the dots, track it for several seconds, and then report their confidence in their tracking as better or worse than their average. In Experiment 1, we manipulated task difficulty with two methods: varying the size of the dot cloud and varying the stability of the target’s velocity. In Experiment 2, the stimulus statistics were fixed and duration of the stimulus presentation was varied. We found similar levels of metacognitive sensitivity in all experiments, with the temporal analysis revealing a recency effect, where error later in the trial had a greater influence on the sensorimotor confidence. In sum, these results indicate humans predominantly monitor their tracking performance, albeit inefficiently, to judge sensorimotor confidence.

**Highlights:**

- Participants consciously reflected on their tracking performance with some accuracy
- Sensorimotor confidence was influenced by recent errors
- Expectations of task difficulty did not play a large role in sensorimotor confidence
- Metacognitive sensitivity of binary confidence judgements on continuous performance can be quantified with standard non-parametric techniques

## 1 Introduction

Sensorimotor decision-making is fundamental for humans and animals when interacting with their environment. For example, it will determine where we look, how we move our limbs through space, or what actions we select to intercept or avoid objects. In return, we may receive decision feedback from the environment, such as resources, knowledge, social standing, injury, or embarrassment. Such outcomes of an action are often crucial for determining subsequent sensorimotor decision-making, particularly in dynamic scenarios where a series of actions are chained together to achieve a sensorimotor goal (e.g., dancing or tracking a target). But what happens if external feedback is absent, partial, or significantly delayed? How then do we judge if an action has been performed well? One possible solution is for the action taker to form their own subjective evaluation of sensorimotor performance using whatever sensory or motor signals are available. These metacognitive judgements reflect the person’s confidence that their action or series of actions accomplished their sensorimotor goal. Yet, despite such judgements being a familiar and everyday occurrence, they have received relatively little direct scientific scrutiny.

Different related elements of sensorimotor confidence have been touched upon in a variety of domains, highlighting the sophisticated monitoring and control processes of the brain that operate on internally-gathered information (see Figure 1 for a summary). At a broader level, there is the topic cognitive control, which describes how goals or plans translate into actual behaviour. It is thought that cognitive control, also known as executive control, is responsible for the appropriate deployment of attention, as well as voluntary selection, initiation, switching, or termination of tasks (Norman and Shallice, 1986; Botvinick et al., 2001; Alexander and Brown, 2010). At a finer level is the study of sensorimotor control. Usually, research questions focus on how the brain senses discrepancies between the intended outcome of motor commands, as specified by an internal model, and the actual action outcomes, that are processed as a feedback signal, to correct and update subsequent motor control signals (Wolpert et al., 1995; Todorov, 2004). While the understanding of sensorimotor processes is quite advanced, both at the behavioural and neural levels, very little is known about our ability to consciously monitor sensorimotor performance.

**Figure 1:**
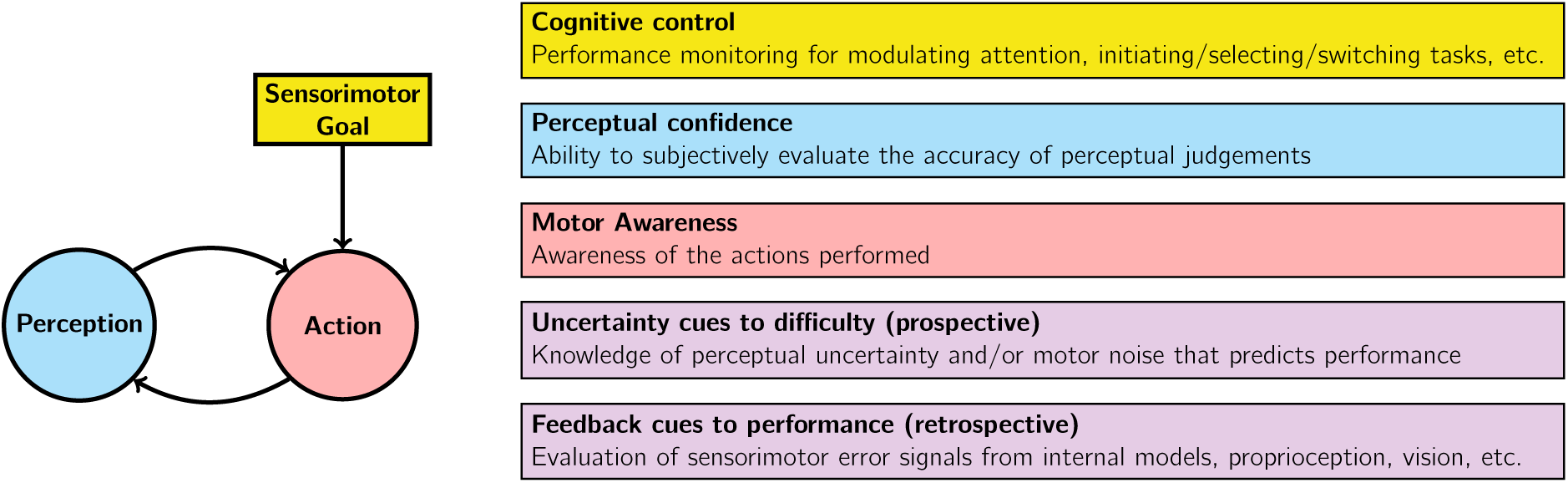
Components of sensorimotor control (left) and related topics in the literature (right). Sensorimotor confidence is a subjective evaluation of how well behaviour fulfilled the sensorimotor goal, considering both sensory and motor factors. The topic of sensorimotor confidence is complementary to the discussions of cognitive control, perceptual confidence, motor awareness, uncertainty, and self-generated feedback. It is likely that cues to difficulty and performance, that are responsible for the computation of sensorimotor confidence, originate both from sensory and motor sources. The former cues are prospective as they are related to how well the acting agent can potentially perform, whereas the latter are retrospective, they become available only after the action has occurred.

If the action is reduced to a simple report of what is perceived, the monitoring of sensorimotor performance reduces to the study of perceptual confidence (Pleskac and Busemeyer, 2010; Fleming and Dolan, 2012; Mamassian, 2016). Perceptual confidence is a metacognitive process, which corresponds to the subjective sense of the correctness of our perceptual decisions (Galvin et al., 2003; Pouget et al., 2016). Human observers exhibit considerable sensitivity to the quality of the processing of sensory information and the resulting ability to predict the correctness of the perceptual choice (Barthelmé and Mamassian, 2010; Kiani et al., 2014; Adler and Ma, 2018). However this so-called Type-2 judgement often incurs additional noise, on top of the sensory noise that reduces perceptual performance (Type-1 decisions) (Maniscalco and Lau, 2016). More recently, researchers have considered the contribution of motor factors in perceptual confidence (Yeung and Summerfield, 2012; Kiani et al., 2014; Fleming and Daw, 2017). Such elements are crucial, for example, for the observer to respond “low confidence” on lapse trials where they mistakenly pressed the wrong key. In other examples, motor behaviour is used as an index of perceptual confidence by tracking hand kinematics while observers report their perceptual judgement (Resulaj et al., 2009; Patel et al., 2012; Dotan et al., 2018). However, these noted contributions are often restricted to simple motor behaviours, and do not take into account motor sources of variability.

Motor awareness, the degree to which we are conscious of the actions we take (Blakemore et al., 2002; Blakemore and Frith, 2003), is also likely to contribute to sensorimotor confidence. Not all actions are consciously monitored, and it is a common experience to act without conscious control. For example, when we are walking, we are not always thinking of exactly how to place one foot in front of the other. Yet, for other actions, we must consciously attend to them, such as threading a sewing needle. A seminal study on motor awareness by Fourneret and Jeannerod (1998) found poor introspective ability for the action made when an unseen hand movement is perturbed by a horizontal displacement in the visual feedback signal. Participants discount their compensatory actions and instead indicated that their hand position followed a trajectory much like the perturbed cursor. Follow-up studies have modified the response to be a binary motor-awareness decision (e.g., “Was feedback perturbed or not”) followed by a confidence rating (Sinanaj et al., 2015; Bègue et al., 2018). Another motor-awareness study measured confidence ratings following a judgement of whether a visual dot was flashed ahead or behind their finger position during up-down movement (Charles et al., 2020). However, we shall argue that none of these measurements of confidence correspond to sensorimotor confidence as we have defined it. Motor-awareness confidence reflects the knowledge held about the executed actions, whereas we propose that sensorimotor confidence involves an additional step of evaluating how well these behaviours serve the sensorimotor goal. For example, a competitive swimmer might know all the moves they executed to reach the other end of the pool (i.e., motor awareness), and confidently report said awareness, but they might find it difficult to judge if they completed this lap faster than usual (i.e., sensorimotor confidence). To our knowledge, the only study to ask participants to explicitly reflect on their sensorimotor performance was by Mole et al. (2018), who had participants perform a virtual driving task. Green lines were placed on the road to indicate a good-performance zone, and after completing the trial, they were asked to report the percentage of time they spent in the green zone (i.e., a continuous measure of sensorimotor confidence). They found that correspondence between objective performance and sensorimotor confidence roughly followed difficulty of the task but was otherwise limited.

The study of sensorimotor confidence should be contrasted with the mere knowledge of sensorimotor uncertainty (Augustyn and Rosenbaum, 2005). In theory, this can be studied by examining how knowledge of variability from sensory, motor, and task sources, influences motor decision-making (Wolpert and Landy, 2012). The majority of studies support the hypothesis that humans plan actions consistent with accurate knowledge of their sensorimotor uncertainty (e.g., Augustyn and Rosenbaum, 2005; Trommershäuser et al., 2008; Stevenson et al., 2009; Bonnen et al., 2015), with some exceptions (e.g., Zhang et al., 2013). However, the degree to which this knowledge is consciously available to the person is highly debatable (Augustyn and Rosenbaum, 2005). Furthermore, judgements of one’s uncertainty in a planned action only allow one to predict the probability of a successful outcome. In this sense, they can act as prospective confidence judgements before the action is taken, but do not constitute retrospective confidence judgements made by reflecting on sensorimotor behaviour from performance monitoring. For example, one would typically have more prospective confidence for riding a bicycle than a unicycle. This belief is not derived from performance monitoring but rather from experience-informed expectation. In other areas of metacognitive research, such use of uncertainty information or other predictions of task difficulty are considered heuristics that can even impair the relationship between objective performance and confidence (e.g., Spence et al., 2015; De Gardelle and Mamassian, 2015; Mole et al., 2018; Charles et al., 2020). Thus, it is desirable to identify the degree to which sensorimotor confidence is based on conscious monitoring of performance from feedback cues versus prospective judgements of performance based on uncertainty cues.

Here, we report on two experiments explicitly measuring sensorimotor confidence in a visuomotor tracking task. In both experiments, participants manually tracked a target, the location of which was inferred from limited noisy sensory information, a twinkling dot cloud, as it followed an unpredictable trajectory. Afterwards, they reported their sensorimotor confidence by subjectively evaluating their performance with a relative judgement of “better” or “worse” than their average. A dynamic task was selected to mirror the sensorimotor goals typically encountered in the real world. In Experiment 1, we manipulated the difficulty of the task by changing the uncertainty of the location of the sensory stimulus with two separate methods. In Experiment 2, we manipulated the stimulus-presentation duration to introduce uncertainty about when the confidence response would be required. We had several goals in this study: 1) to test whether humans are able to make reasonable sensorimotor confidence judgements from monitoring performance error signals rather than relying only on uncertainty-based expectations; 2) to quantify how well sensorimotor confidence reflected objective performance; and 3) to examine what evidence contributed to the sensorimotor confidence judgement.

## 2 Experiment 1

Experiment 1 sought to measure sensorimotor confidence in a visuomotor tracking task and establish a metric of metacognitive sensitivity that quantified how well the confidence judgements corresponded to objective tracking performance. Difficulty in the task was manipulated in the *cloud-size* session by varying the external noise of the sensory evidence indicating the target location. In the *velocity-stability* session, we varied the degree of noise in the target’s trajectory. To investigate the error evidence contributing to the sensorimotor confidence, we investigated the temporal pattern of metacognitive sensitivity, applying our metric to 1 s time bins within the trial.

### 2.1 Methods

#### Participants

Seven naive participants (23 – 35 years old, one left-handed, one female) took part in the study. All had normal or corrected-to-normal vision and self-reported normal motor functioning. They received details of the experimental procedures and gave informed consent prior to the experiment. Participants were tested in accordance with the ethics requirements of the École Normale Supérieure and the Declaration of Helsinki.

#### Apparatus

Stimuli were displayed on a V3D245 LCD monitor (Viewsonic, Brea, CA; 52 × 29.5 cm, 1920 × 1080 pixels, 60 Hz). Participants sat 46.5 cm from the monitor with their head stabilised by a chin rest. Manual tracking was performed using a Logitech M325 wireless optical mouse, operated by the participant’s right hand. Subjective assessments of performance were reported on a standard computer keyboard with the left hand. The experiment was conducted using custom-written code in MATLAB version R2014a (The MathWorks, Natick, MA), using Psychtoolbox version 3.0.12 (Brainard, 1997; Pelli, 1997; Kleiner et al., 2007).

#### Dot-cloud stimulus

Every frame, two white dots were drawn from a 2D circularly symmetric Gaussian generating distribution with standard deviation *σ*_cloud_. The mean of the distribution was the tracking target, which was invisible to observers and must be inferred from the dot cloud. Each dot had a one-frame lifetime and new dots were drawn every frame. Due to the persistence of vision, participants had the impression of seeing up to 10 dots at any one time (Figure 2A). Dots had a diameter of 0.25 deg and were presented on a mid-grey background. Dots were generated using Psychtoolbox functions that rendered them with sub-pixel dot placement and high quality anti-aliasing. The horizontal position of the target changed every frame according to random walk in velocity space (Figure 2B): *v*_*t*+1_ = *v*_*t*_ + *ϵ* and *ϵ* ∼ 𝒩 (0, *σ*_walk_) deg/s. This gave the target momentum, making it more akin to a real-world moving target (Figure 2C). Both the target and the black cursor dot (diam.: 0.19 deg) were always centred vertically on the screen. Trajectories that caused the target to move closer than 2 × max(*σ*_cloud_) from the screen edge were discarded and resampled prior to presentation.

**Figure 2:**
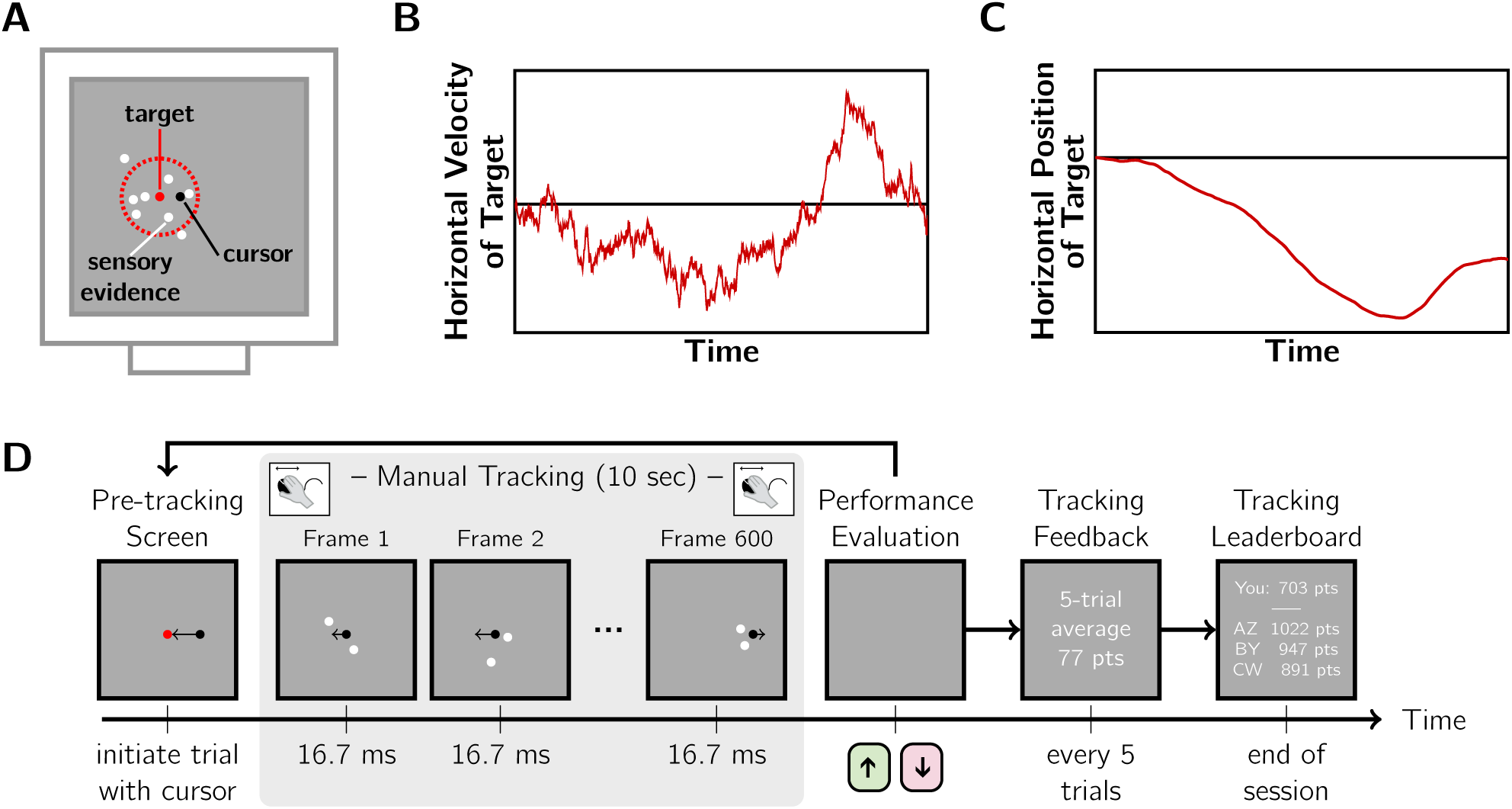
Visuomotor tracking task. A: The “twinkling” dot cloud stimulus (white), generated by drawing two dots per frame from a 2D Gaussian generating distribution. Red: mean and 1 SD circle, which were not displayed. Black: mouse cursor. The dots provided sensory evidence of target location (generating distribution mean). As shown, more than two dots were perceived at any moment due to temporal averaging in the visual system. B: Example target random-walk trajectory in velocity space. C: The corresponding horizontal trajectory of the target. D: Trial sequence. Trials were initiated by the observer, followed by 10 s of manual tracking of the inferred target with a computer mouse. Then, participants reported their sensorimotor confidence by indicating whether their performance on that trial was better or worse than their average. Objective performance feedback was provided intermittently including average points awarded and a final leaderboard. Difficulty manipulations: cloud size (*σ*_cloud_) and velocity stability (*σ*_walk_) were varied in separate sessions.

#### Task

The trial sequence (Figure 2D) began with a red dot at the centre of the screen. Participants initiated the tracking portion of the trial by moving the black cursor dot to this red dot, causing the red dot to disappear. The dot-cloud stimulus appeared immediately, with the target centred horizontally. The target followed its horizontal random walk for 10 s. Then, the participant made a subjective assessment of tracking performance while viewing a blank grey screen, reporting by keypress whether they believed their tracking performance was better or worse than their session average. The experiment was conducted in two sessions on separate days. In the “cloud size” session, the standard deviation of the dot cloud, *σ*_cloud_, was varied from trial to trial (5 levels: 1, 1.5, 2, 2.5, and 3 deg) and the standard deviation of the random walk, *σ*_walk_, was fixed at 0.15 deg/s. In the “velocity stability” session, *σ*_walk_ was varied (5 levels: 0.05, 0.10, 0.15, 0.20, and 0.25 deg/s) and *σ*_cloud_ was fixed at 2 deg. Examples of the stimuli for both sessions are provided as Supplementary media files. The order of sessions was counterbalanced across participants to the best extent possible. Each session began with a training block (20 trials, 4 per stimulus level in random order), where only tracking responses were required. The training trials allowed participants to become familiar with the stimulus and set-up, and to form an estimate of their average performance. The main testing session followed (250 trials, 50 per stimulus level in random order). For the second session, participants were instructed to form a new estimate of average performance, and not to rely on their previous estimate.

#### Grading objective performance

For all analyses, we used root-mean-squared-error (RMSE) as our measure of tracking error, calculated from the horizontal distance between the target (i.e., the current distribution mean) and the cursor. For the purposes of feedback, the tracking performance on each trial was converted to a score according to the formula *points* = 100 − 30 ∗ *RMSE*. Typical scores ranged from 60 to 80 points. Every 5 trials, the average score for the previous 5-trials was reported. This feedback was provided for both training and test trials. Presenting the average score served several purposes. It 1) incentivised good tracking behaviour; 2) encouraged consistent performance across the session; and 3) helped participants to maintain a calibrated internal estimate of average performance. At the end of a session, participants were shown their cumulative score for that session and ranking on a performance leaderboard.

#### Metacognitive sensitivity metric

To examine sensorimotor confidence, we sought a metacognitive sensitivity metric that reflected how well the confidence reports identified good from bad tracking performance (i.e., low versus high RMSE). This concept is similar to the one used in perceptual confidence, where metacognitive sensitivity refers to a person’s ability to distinguish correct from incorrect decisions (Fleming and Lau, 2014). As the outcome of tracking was not binary (e.g., correct vs. incorrect), we considered the objective tracking performance within a trial relative to all trials within the session performed by that participant. We constructed two objective-performance probability distributions conditioned on the sensorimotor confidence: one distribution for trials followed by a “better than average” response and one for “worse than average” responses (Figure 3A-B). A high overlap in these conditional distributions would reflect low metacognitive sensitivity as this means objective performance is a poor predictor of the participant’s evaluation of their performance. Conversely, low overlap indicates high metacognitive sensitivity. We used an empirical Receiver Operating Characteristic (ROC) curve, also known as a quantile-quantile plot (Figure 3C), for a non-parametric measure of metacognitive sensitivity that reflected the separation of these distributions, independent of any specific criterion for average performance. As shown in Figure 3D, completely overlapping distributions would fall along the equality line in an ROC plot, resulting in an Area Under the ROC curve (AUROC) of 0.5. In contrast, complete separation would yield an AUROC of 1. An advantage of this technique over methods that rely on averaging (e.g., classification images) is that this method is suitable for continuous performance distributions of any shape (e.g., skewed). There are two things worth noting about the interpretation of this metric. First, this is not the ROC method other researchers typically use to measure perceptual confidence (Barrett et al., 2013; Fleming and Lau, 2014). AUROC has, however, been used previously to explore the relationship between choice correctness and continuous confidence ratings as well as reaction times (Faivre et al., 2018). Second, our AUROC measure has the following interpretation: if the experimenter was given the RMSE of two trials and was told one was rated “worse” and the other as “better”, the AUROC would reflect the probability of correctly inferring that the objectively better trial of the two was rated as “better” by the participant.

**Figure 3:**
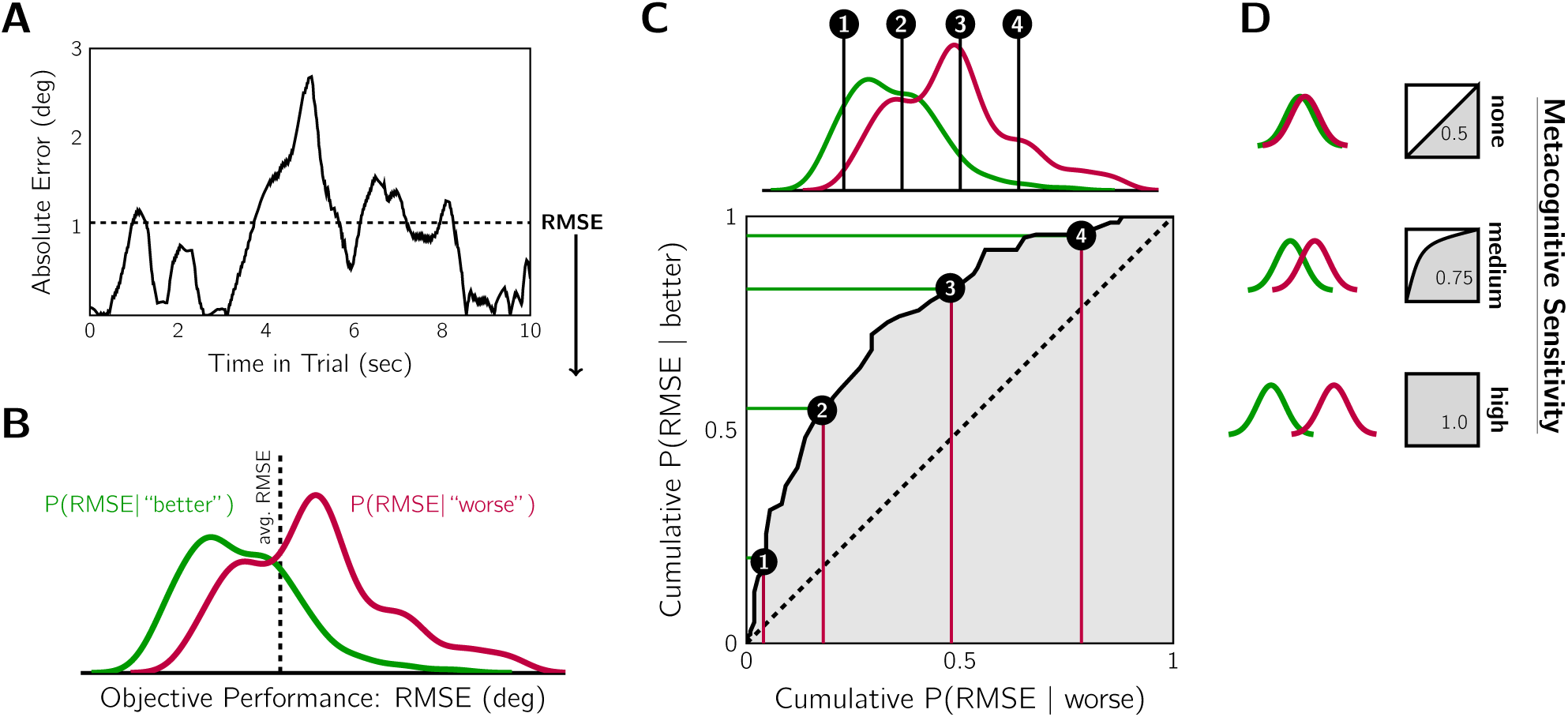
A metacognitive sensitivity metric. A: Example of tracking error within a trial. Root-mean-squared-error (RMSE, dashed line) was the objective performance measure. B: Example participant’s objective-error distributions, conditioned on sensorimotor confidence, for all trials in the variable cloud-size session. True average performance (dashed line) indicates the ideal criterion. Smaller RMSE tended to elicit “better” reports, and larger RMSE “worse”. C: Metacognitive sensitivity was quantified by the separation of the conditional objective-error distributions with a non-parametric calculation of the Area Under the ROC (AUROC) using a quantile-quantile plot. At every point along the objective-performance axis, the cumulative probability of each conditional error distribution was contrasted. D: The area under the resulting curve is the AUROC statistic, with 0.5 indicating no meta-cognitive sensitivity and 1 indicating maximum sensitivity. The greater the separation of the conditional distributions, the more the objective tracking performance was predictive of sensorimotor confidence, and thus the higher the metacognitive sensitivity.

### 2.2 Results

#### Confirming the difficulty manipulation

We first examined whether the difficulty manipulation affected objective tracking performance. Figure 4A shows the mean RMSE for each stimulus level for the two difficulty manipulations. Qualitatively, the difficulty levels appear matched for most participants: performance curves follow the equality line. To check this result, we fit a generalised linear mixed-effects model (GLMM) to the RMSE values of each trial. The fixed effects in the model were difficulty manipulation (cloud-size or velocity-stability), stimulus difficulty (five levels), trial number, and an intercept term. The random effect was the participant. Trial number was included to test whether learning occurred during the experiment. An analysis of deviance was performed using Type II Wald chi-square tests, revealing that only difficulty level had a significant effect on tracking performance (*χ*^2^ = 2042.85, *p* < .001). This confirms that the difficulty manipulations had an impact on tracking performance. Also, the effect appears to be comparable across the difficulty manipulations, as neither difficulty manipulation nor an interaction between difficulty manipulation and stimulus level had a significant effect (*p* > 0.05). As trial number did not have a significant effect either (*p* > 0.05), training trials were likely sufficient for performance to stabilise prior to the main task.

**Figure 4:**
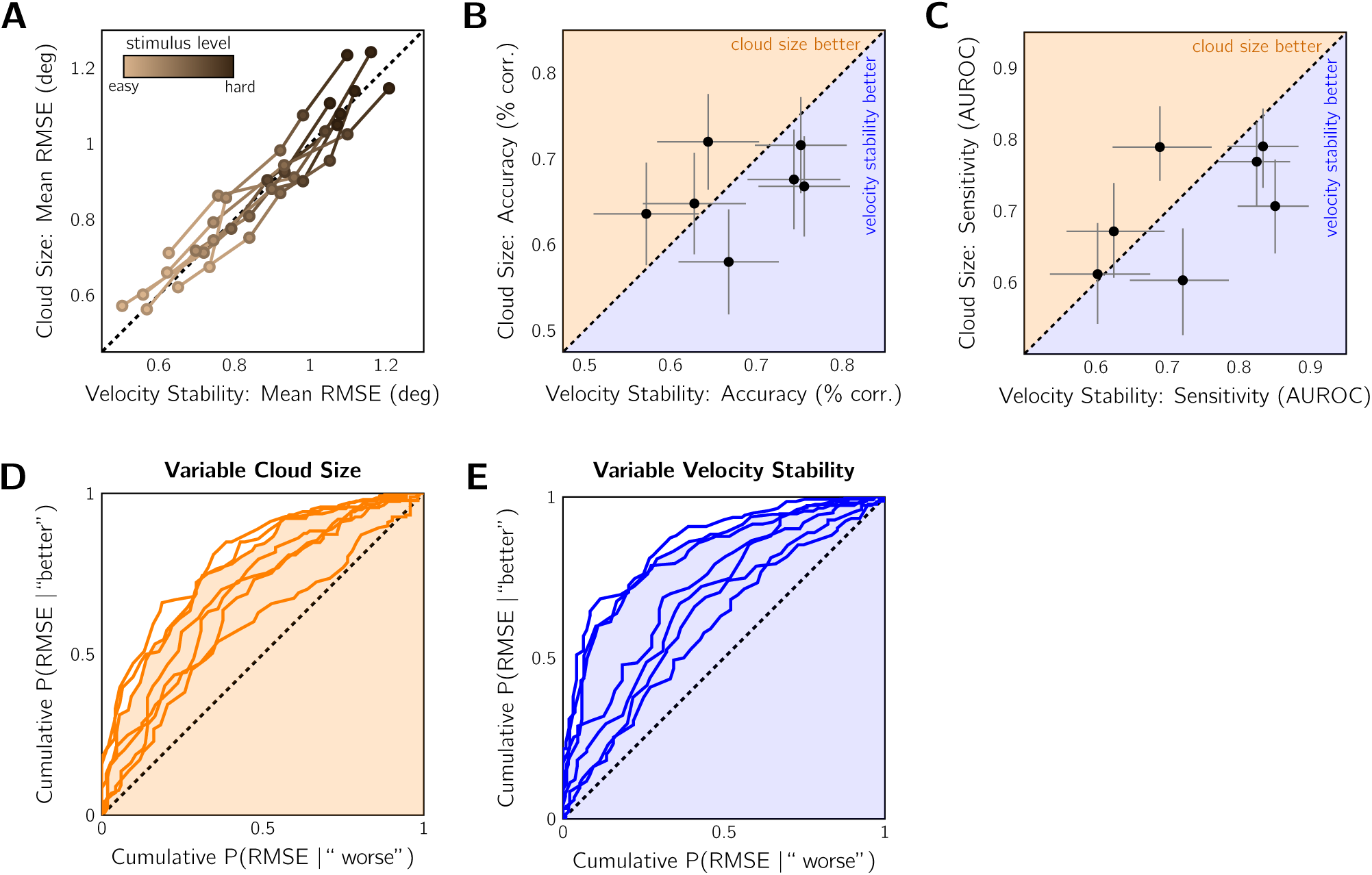
Comparable above-chance metacognitive sensitivity for cloud-size and velocity-stability difficulty manipulations in Experiment 1 (*n* = 7). A: Effect of difficulty manipulation on tracking error. Mean RMSE contrasted for equivalent difficulty levels in the variable cloud-size session and the variable velocity-stability session. Colour: difficulty level. Curves: individual participants. Dashed line: equivalent difficulty. B: Comparison of metacognitive accuracy for the two difficulty-manipulation techniques, pooled across difficulty levels. Data points: individual subjects. Dashed line: equivalent accuracy. Error bars: 95% binomial SE. Shaded regions indicate whether metacognitive accuracy was better for the cloud-size or velocity-stability session. C: Same as in (B) but comparing the sensitivity of the sensorimotor confidence judgement. Dashed line: equivalent sensitivity. Error bars: 95% confidence intervals by non-parametric bootstrap. D: ROC-style curves for individual participants in the cloud-size session, pooled across difficulty levels. Shading: AUROC of example observer. Dashed line: the no-sensitivity lower bound. E: Same as (D) for the velocity-stability session. Shading corresponds to the same example observer.

### Overall metacognitive accuracy

Next, we examine metacognitive accuracy, which is the percentage of trials correctly judged as better or worse than average. Performance in both sessions was significantly better than chance (cloud-size session: 66.3 ± 1.8% correct; velocity-stability session: 68.0 ± 2.7%). The accuracy results for each session are contrasted in Figure 4B. Two participants had significantly higher accuracy in the cloud-size session, according to the 95% binomial error confidence intervals, and three participants were significantly more accurate in the velocity-stability session. Overall, evaluation of tracking performance was similar in the two conditions. However, this accuracy metric may be subject to response bias. Therefore, we examined meta-cognitive sensitivity.

### Overall metacognitive sensitivity

A similar pattern of results was found for metacognitive sensitivity. Metacognitive sensitivity (AUROC, see Methods) is contrasted between the sessions in Figure 4C and the individual ROC-style curves for cloud-size session and the velocity-stability session are shown in Figures 4D and 4E, respectively. All participants displayed some degree of metacognitive sensitivity in both sessions: none of the ROC-style curves fell along the equality line. On average, the AUROC in the cloud-size session was 0.71 ± 0.03 (mean±SEM) and was 0.74 ± 0.04 for the velocity-stability session. At the group level, a Wilcoxon’s Matched-Pairs Signed-Ranks Test revealed no significant difference between AUROCs from the two sessions (*n* = 7, *T* = 7, *p* > 0.05). To examine the sensitivity at the individual subject level, we performed a bootstrap procedure in which the AUROC was computed for each participant 1000 times, sampling from their trial set with replacement, allowing us to calculate 95% confidence intervals for our estimates (Figure 4C). Three participants were significantly more sensitive in the velocity-stability session, one was significantly more sensitive in the cloud-size session, and the other three showed no significant difference between the two conditions. It is unlikely that these results are due to a learning effect across sessions: 3 of the 4 significant results come from greater meta-cognitive accuracy in the first session completed. Another consideration is the amount of variability in performance for each individual and session. A highly variable participant may have a higher metacognitive sensitivity score because distinguishing better from worse performance is easier if a better trial differs more, on average, from a worse trial. Also, variance could have differed between the two difficulty manipulations, affecting within-participant comparisons of metacognitive sensitivity. To examine this we fit a GLMM of the AUROC with participant as the random effect, and fixed effects of RMSE variance (pooled across difficulty levels), difficulty manipulation, and an intercept term. We found no significant effect of any of our predictors, indicating that performance variance likely did not play an important role in determining metacognitive sensitivity.

### Temporal profile of metacognitive sensitivity

We conducted an analysis of metacognitive sensitivity for each 1 s time bin within the 10 s trial to examine the degree to which each second of tracking contributed to the final sensorimotor confidence judgement. An AUROC of 0.5 indicates that error in that 1 s time bin has no predictive power for the metacognitive judgement; an AUROC of 1 indicates perfect predictive power. Figure 5A shows the results of this analysis. In both the cloud-size and the velocity-stability sessions there was a noticeable recency effect: error late in the trial was more predictive of sensorimotor confidence than error early in the trial. There was no discernible difference between the two difficulty manipulations, except for the first few seconds where early error was more predictive for the velocity-stability session.

**Figure 5:**
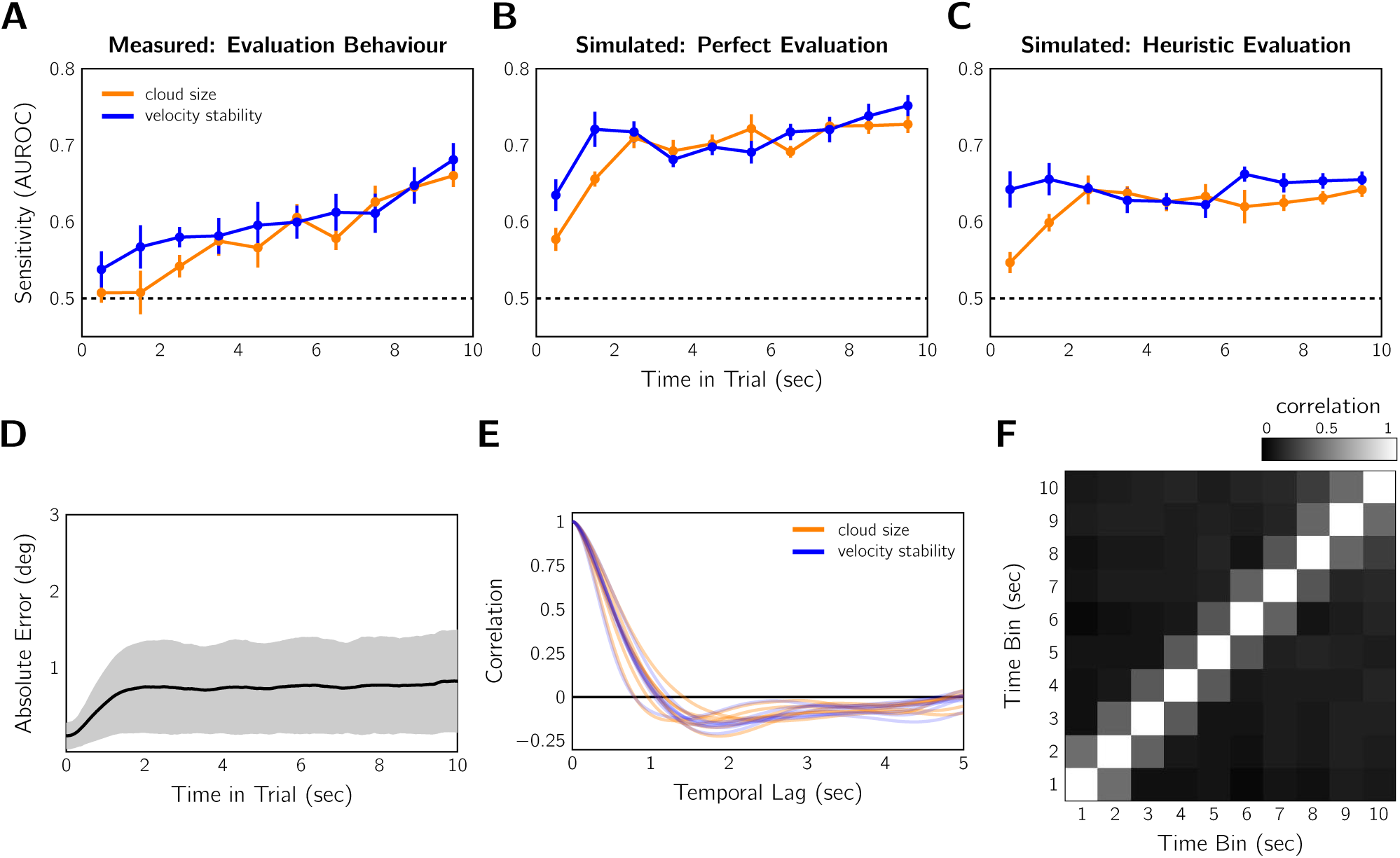
Performance weighting over time for sensorimotor confidence in Experiment 1 (*n* = 7). A: AUROC analysis performed based on each 1-s time bin in the tracking period. Error bars: SEM across participants. Error later in the trial is more predictive of sensorimotor confidence as indicated by the higher AUROC. B: The same analysis as in (A) for an ideal observer that has perfect knowledge of the error and compares the RMSE to the average RMSE. C: Temporal analysis performed with simulated responses based on expected performance according to the heuristic of difficulty level (see text). D: Mean absolute error between target and cursor for all trials in both sessions shows error plateaus between 1-2 s and remains stable for the remainder of the trial. Grey region: ±1 SD. E: Auto-correlation of the tracking error signal for each subject and each session. F: Autocorrelation matrix of the 1 s binned RMSE. Data pooled over trials, conditions, and participants. The correlation between time-bins is relatively low after 1 s.

For comparison, we also computed the temporal AUROCs, replacing the participant’s responses with simulated sensorimotor confidence judgements under two strategy extremes. Figure 5B shows the AUROC time course for an ideal observer that had perfect knowledge of performance (RMSE) and based the confidence judgment on whether the RMSE was truly better or worse than average (i.e., weighted all time points equally). After the first two seconds of tracking, the temporal AUROC is relatively level. Note that no time bin was perfectly predictive of the confidence judgement, because the error within one second is not equivalent to the total error across the entire trial. Figure 5C shows the AUROC time course for an observer that perfectly uses uncertainty cues to judge the difficulty level of the trial, and computes prospective confidence rather than basing the confidence judgment on performance monitoring. Again, no single time bin should be particularly informative if one is assessing a cue (e.g., dot-cloud size, velocity stability, etc) that does not disproportionately occur at or affect performance for one particular portion of the trial. Confidence was coded as “worse” for the two hardest difficulty levels, “better” for the two easiest, and flipping a 50-50 coin for the middle difficulty level. Again, both temporal profiles are flat after the first 2 s. Neither perfect monitoring nor prospective confidence based on uncertainty cues produced the recency effect in measured metacognitive behaviour. This result, however, is not trivial due to the complex correlation structure of the error signal, which we investigated next.

Weighing all time points equally is only an optimal strategy if all time bins are equally predictive of trial-averaged performance. Error variability is one factor that can affect that: periods of low error volatility have less impact on the predictive validity of a time bin for overall RMSE. Thus, a recency effect might be an optimal strategy if there is higher error volatility late in the trial. We found that error is overall lower and less variable before 2 s (Figure 5D). This is because participants begin the trial by placing their cursor at the centre of the screen, where the target is located. After this initial 2 s, however, tracking error is constant in both mean and variance, indicating that all these time points are equally informative on average about the final RMSE. Thus, error variance may explain why metacognitive sensitivity was reduced for the initial 2 s for the measured and simulated sensorimotor confidence, but it cannot explain the observed recency effect. Figure 5E shows the auto-correlation of the signed error signal for each participant averaged across difficulty levels. This graph reveals that error is correlated up to ±1 s, and is slightly anti-correlated thereafter. Errors are necessarily related from moment to moment, due to the continuous nature of tracking. To resolve a tracking error, one needs to make a corrective action to compensate. The anti-correlation is likely a result of such corrective actions. Figure 5F shows that this salient auto-correlation up to ±1 s is also present between the RMSE of neighbouring 1 s time bins. These results indicate that some of the predictive power of error in one time bin may be attributed to weighting of error in a neighbouring bin. Thus, if we ask for what *additional* variance is accounted for, starting with last bin, the recency effect would appear even stronger.

### Other performance metrics

Our modelling thus far has been based on the error between the location of the target and the cursor placement. However, this is not a realistic model of how the participant perceives their error as they imperfectly infer target location from the dot cloud, which is predominately affected by the external noise *σ*_cloud_. To model this perceptual process (Figure 6A), we opted for a simple exponential filtering of the centroid signal (i.e., the mid-point of the two dots presented each frame). The true centroid position is a reasonable input, given that humans perform well at static centroid estimation (McGowan et al., 1998; Juni et al., 2010). The smoothing aims to capture both the temporal averaging in the visual system, which causes a cloud of 10 or so dots to be perceived, as well as the averaging across time for strategic decision-making (Kleinman, 1969; Bonnen et al., 2015). The current estimate of target position, 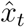 is obtained by computing the weighted average at time, *t*, of the horizontal component of the current centroid, *c*_*t*_, with the previous estimate, 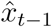:

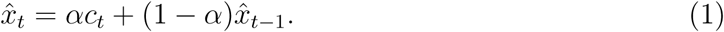

**Figure 6:**
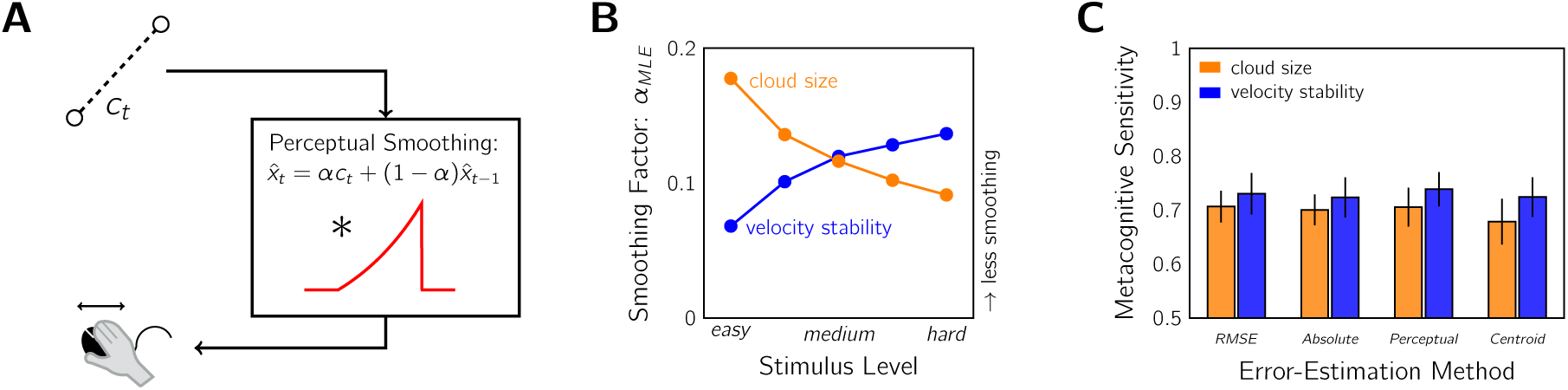
Comparing metacognitive sensitivity with different error-estimation methods. A: Diagram of the exponentially-smoothed perceptual model. Input: horizontal position of dot-cloud centroid, *c*_*t*_ (i.e., dot midpoint on single frame). The perceptual system smooths the signal by convolving with an exponential to produce the target estimate 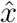. This is equivalent to the weighted sum of current input and previous estimate, 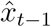, according to the smoothing parameter, *α*. Output: perceived error determines the motor response. B: Setting of *α* that minimises the difference between true and perceived target location for each difficulty level and condition. C: Metacognitive sensitivity AUROC as measured under several error-estimation methods. RMSE: root-mean-squared-error between true target location and cursor. Absolute: mean absolute error between target and cursor. Perceptual: error according to the perceptual model in (A) with *α* values from (B). Centroid: RMSE calculated using dot-cloud centroid rather than true target location.

The smoothing parameter, *α*, controls the steepness of the exponential. Larger *α* mean that current sensory evidence is weighted more than previous target estimates, and vice versa. The weighting is a trade-off that has to be balanced: averaging improves the amount of information contributing to the estimate, but too much averaging into the past leads to biased estimates.

We selected the value of *α* that minimised the sum of squared errors between true target location and the model’s estimate as a stand in for the observer’s estimate of the current location of the target. This was calculated separately for each stimulus level and condition (Figure 6B). As expected, there is less smoothing (larger *α*) for the easy, small dot clouds than the more difficult, large dot clouds (smaller *α*). This is because accepting some history bias only makes sense when dealing with the noisier large dot clouds. The opposite pattern is true for the velocity-stability condition. If velocity stability is high (easy), it is safer to average further into the past to improve the estimate than if velocity stability is low (hard). When the AUROC was calculated from the trial RMSE according to this perceptual model, however, the results are relatively unchanged (Figure 6C). In fact, using the RMSE based on the raw centroid signal also produced a similar AUROC estimate. We also examined the mean absolute error, which is an alternate objective tracking metric to using RMSE, and obtained the same result. The unchanging AUROC across these performance metrics is likely due to the high correlation between all of these error measures. As compared to the RMSE method, the correlations for the cloud-size condition are *r* = 0.98, 0.93, and 0.77 for absolute error, perceptual error, and centroid error respectively. For the velocity-stability condition, these are *r* = 0.98, 0.94, and 0.94. This is because all methods are measures of the mean performance, which will change little with unbiased noise if given sufficient samples (i.e., 10 s of tracking). Thus, we conclude that our AUROC statistic was a robust measure and that the overlap in the confidence-conditioned distributions is unlikely due to the selection of RMSE as the objective-performance metric.

### Summary

Experiment 1 measured sensorimotor confidence for visuomotor tracking, under both cloud-size and velocity-stability manipulations of difficulty, to address the three goals of this study. A robust AUROC statistic, that quantified the ability of the confidence judgements to distinguish objectively good from bad tracking, indicated that confidence judgements were made with comparable above-chance metacognitive sensitivity for both difficulty manipulations. Furthermore, a temporal analysis revealed a recency effect, where tracking error later in the trial was found to disproportionately influence sensorimotor confidence. We propose that this is due to imperfect performance monitoring and not prospective confidence based on heuristic cues to difficulty (i.e., cloud size, velocity stability).

## 3 Experiment 2

The goal of Experiment 2 was to further investigate the recency effect. To this end, we repeated the task keeping the stimulus statistics fixed (*σ*_cloud_ and *σ*_walk_) and instead varied the duration of the stimulus presentation in an interleaved design. This made the time when the sensorimotor-confidence judgement was required less predictable. Thus, participants would be encouraged to sample error evidence for their confidence throughout the trial instead of waiting until the final portion of the stimulus duration. If a response-expectation strategy was the cause of the recency effect, we would expect to see flatter temporal AUROCs for this mixed-duration design. Otherwise, if the recency effect is due to a processing limitation of sensorimotor confidence, we would expect error in the last few seconds to largely determine sensorimotor confidence regardless of the duration condition. Additionally, this experiment allowed us to investigate sensorimotor confidence in the context of a fixed difficulty setting that encourages participants to monitor their performance. This is because prospective judgements of confidence, based on sensorimotor uncertainty, are uninformative when the stimulus statistics are unchanging.

### 3.1 Methods

#### Participants

There were seven new participants in Experiment 2 (21–31 years old, one left-handed, four female). All participants had normal or corrected-to-normal vision and no self-reported motor abnormalities. Participants were naive to the purpose of the studies except one author. Prior to the experiment, the task was described to the participants and consent forms were collected. Participants were tested in accordance with the ethics requirements of the Institutional Review Board at New York University.

#### Apparatus

All experiments were conducted on a Mac LCD monitor (Apple, Cupertino, CA; late 2013 version, 60 × 34 cm, 1920 × 1080 pixels, 60 Hz), with participants seated 57 cm from the monitor. Participants operated a Kensington M01215 wired optical mouse with their right hand when manually tracking the stimulus. Subjective performance evaluations were collected on a standard computer keyboard. Experiments were conducted using custom-written code in MATLAB version R2014a (The MathWorks, Natick, MA), using Psychtoolbox version 3.0.12 (Brainard, 1997; Pelli, 1997; Kleiner et al., 2007).

#### Task

Stimulus presentation duration was manipulated with an interleaved design and three levels (6, 10, and 14 s) while the stimulus statistics remained fixed at *σ*_cloud_ = 2 deg and *σ*_walk_ = 0.15 deg/s. Data were collected over three sessions, with each session composed of 15 training trials (5 per duration, randomised order) followed by 225 test trials (75 per duration, randomised order). Again, after each stimulus presentation, participants rated their subjective sense of their tracking performance as either “better” or “worse” than their session average. As shown in Experiment 1, tracking before 2 s in this task has a different error profile, due to the target and cursor both starting at the same location from stationary. We opted to not count these initial 2 s of tracking in the final score so that trial duration could not serve as a difficulty manipulator in this experiment (e.g., a 6 s trial is more likely to have lower RMSE than a 14 s trial). In order to signal when the tracking contributed to the final score, the cursor was initially red (not contributing) and switched to green (contributing to the score) after 2 s. Furthermore, to ensure that all trials had the same stimulus statistics, all trajectories were sampled based on a 14 s stimulus and accepted or rejected before being temporally truncated if necessary. Tracking performance was scored and feedback given in the same manner as the previous experiment.

### 3.2 Results

In Experiment 2, we manipulated the duration of stimulus presentation with three interleaved conditions of 6, 10, or 14 s. The consequence of duration on objective tracking performance was a small decrease in performance with longer durations (Figure 7A). The sensorimotor confidence judgements also showed slightly lower metacognitive accuracy (Figure 7B) and sensitivity (Figure 7C) for longer durations. Overall, the average AUROC from pooling data across durations was 0.68 ± 0.04 SEM (Figure 7D) and all participants had above-chance metacognitive sensitivity according to bootstrapped confidence intervals calculated as per the same procedure as Experiment 1. When split by session, the AUROCs were 0.68 ± 0.04, 0.68 ± 0.03, and 0.71 ± 0.02, suggesting that metacognitive performance was relatively unchanging across the sessions. Note that for these analyses we discarded the initial 2 s of tracking that the participants were instructed to ignore.

**Figure 7:**
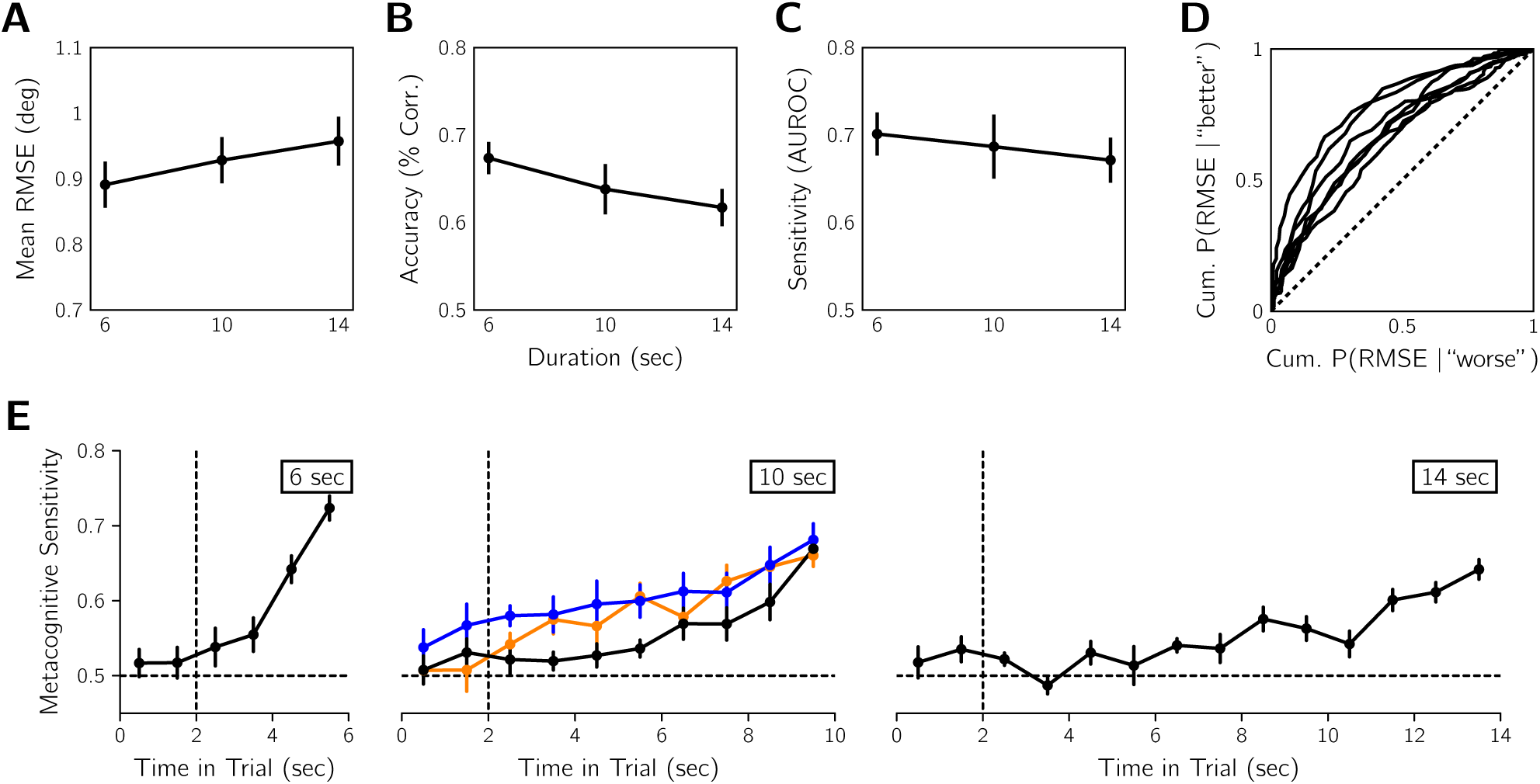
Recency effect replicated for variable stimulus-presentation duration in Experiment 2 (*n* = 7). A: Mean objective tracking performance for each duration condition averaged across observers. B: Sensorimotor-confidence accuracy for each duration condition. C: Metacognitive sensitivity for each duration condition. D: ROC-style curves for individual participants for AUROC pooled across durations. Dashed line: the no-sensitivity lower bound. Error before 2 s was excluded from the calculations in panels A-D. E: Temporal AUROCs calculated for 1 s time bins for each duration condition averaged across participants for Experiment 2 (black). For comparison, the results in Figure 5A are replotted (orange: cloud-size session; blue: velocity-stability session). Vertical dashed line at 2 s indicates the timing of cursor colour-change cue to begin evaluating tracking. Horizontal dashed line the no-sensitivity line. Error bars in all graphs are SEM.

Figure 7E shows the temporal profile of metacognitive sensitivity for each duration as well as the results from Experiment 1. Participants were instructed to ignore tracking error occurring before 2 s, when the cursor changed colour, for estimating sensorimotor confidence, and we observed low metacognitive sensitivity for these time points. Due to RMSE being partially correlated between adjacent time bins (Figure 5F), slightly elevated sensitivity for the time bin at 2 s does not necessarily indicate non-compliance with task instructions. For the remainder of the trial, later time points tend to have higher metacognitive sensitivity, consistent with the recency effect observed in Experiment 1. The steepness of the temporal AUROC was also greater for shorter trial durations. This is to be expected as the contribution of a 1 s time bin to the final RMSE is greater when the trial is short. A recency effect is also consistent with the observed lower overall metacognitive performance for longer durations, because a smaller percentage of the total error signal contributes to sensorimotor confidence.

We observed a steeper recency effect for the 10 s condition than for the cloud-size or velocity-stability sessions in Experiment 1. Specifically, the sensitivity for the final time bin is comparable, but almost all other time bins have lower sensitivity than for the difficulty-manipulation results. However, it is unclear the extent to which the request that participants ignore the first 2 s of tracking contributed to this difference, as this instruction was not given in the previous experiment. We also attempted to compare the temporal AUROCs quantitatively with mixed success (see Supplementary Information). We found evidence for a stronger recency effect for Experiment 2 than Experiment 1. Furthermore, in our supplementary analyses, accounting for the recency effect and/or external noise via our perceptual model in Figure 5A gave little benefit when attempting to predict sensorimotor confidence for either experiment (at most ∼ 5% increase in predictive accuracy). However, we caution against strong conclusions from these supplementary analyses as certain properties of the obtained data set were not ideal for these quantitative model fits.

In sum, We replicated the recency effect of Experiment 1 for all stimulus durations. Thus the final few seconds of tracking had the greatest influence on sensorimotor confidence regardless of whether the participant knew when the stimulus would terminate. This suggests that response-expectation is unlikely to be the source of the recency effect.

## 4 Discussion

In two experiments, participants completed a visuomotor tracking task where trials were followed by a sensorimotor confidence judgement of “better” or “worse” than average tracking performance. We calculated the degree to which these judgements predicted objective tracking for manipulations of task difficulty (Experiment 1) and trial duration (Experiment 2), with an AUROC metacognitive-sensitivity statistic that ranged from no sensitivity at 0.5 and perfect sensitivity at 1. In both experiments we found above-chance metacognitive sensitivity and a temporal profile that suggested that error later in the trial contributed more to sensorimotor confidence.

### 4.1 Performance monitoring

One of our aims was to establish if humans would actively monitor their own performance to judge sensorimotor confidence. An alternate strategy would be to use cues to uncertainty (e.g., cloud size) to predict task difficulty and thus the likelihood of performing well. From our experiments, we found several indicators of performance monitoring. First, Experiment 1 manipulated task difficulty systematically with two methods, varying either the cloud-size parameter (*σ*_cloud_) or the velocity stability parameter (*σ*_walk_). The manipulation of *σ*_cloud_ was very noticeable, with all participants reporting the stimulus manipulation in their debriefing interviews, whereas varying *σ*_walk_ was more subtle and participants had difficulty identifying the manipulation (supplementary media files are provided to demonstrate the difficulty manipulations). Thus, if the strategy was to rely exclusively on cues to uncertainty, and given that the manipulations had sizeable and comparable effects on tracking performance, we would expect higher metacognitive sensitivity for the cloud-size session than the velocity-stability session. We did not find supporting evidence as there was no significant difference in sensitivity between the sessions.

Stronger supporting evidence was found in Experiment 2, where task difficulty was kept the same for all trials by fixing the stimulus statistics. In this scenario, there are no explicit uncertainty cues for the participant to use. Yet, metacognitive sensitivity was only slightly lower than was observed in Experiment 1 (AUROC of 0.68 in Experiment 2 versus 0.71 for cloud-size and 0.74 for velocity-stability in Experiment 1). However, variability in tracking performance is not the same for fixed- and variable-difficulty designs; RMSE differences are likely to be lower for a fixed-difficulty design, complicating the comparison. Furthermore, the difficulty manipulation in Experiment 1 may have permitted a mixed strategy, combining performance monitoring and uncertainty heuristics. Thus, our results from Experiment 2 supporting the performance-monitoring hypothesis are a better indicator of how well performance monitoring captures true tracking performance than the results of Experiment 1.

The best evidence for performance monitoring is the recency effect we observed in both experiments. This is because it demonstrates that some moments in the trial influences sen-sorimotor confidence more than others. Such a result is unlikely from the use of uncertainty cues. For the cloud-size session, all time points equally signal the uncertainty, so there is no reason that the final seconds should be privileged. Similarly, for the velocity-stability session, the behaviour of the target would have to be observed for some period of time to assess velocity stability, but this could be done at any point during the trial. One possibility is that participants were waiting until the end of the trial to make these assessments, but the results of Experiment 2 argue against this, as the recency effect was still found when stimulus-presentation duration was randomised. If instead participants were using some other heuristic strategy (e.g., average velocity, amount of leftward motion, etc.), this would also not produce a recency effect unless it predicted performance later in the trial but not early performance. From an information-processing standpoint, performance monitoring is likely to exhibit temporal sub-optimalities due to either leaky accumulation of the error signal during tracking (Busemeyer and Townsend, 1993; Smith and Ratcliff, 2004) or the temporal limitations of memory for retrospective judgements (Atkinson and Shiffrin, 1968; Davelaar et al., 2005).

Before we examine the recency effect, we first comment on the possibility of a mixed strategy of performance monitoring and uncertainty heuristics. Metacognitive judgements based on a mixed strategy combining actual performance and cues to uncertainty have been reported for sensorimotor confidence (Mole et al., 2018), motor-awareness confidence (Charles et al., 2020), and perceptual confidence (De Gardelle and Mamassian, 2015; Spence et al., 2015), with some exceptions (e.g., Barthelmé and Mamassian, 2010). Yet, it is unclear if a mixed strategy was used in Experiment 1 of the present study. The obvious/subtle distinction between the cloud-size and velocity-stability sessions that favours a purely performance-monitoring strategy was not supported by a rigorous empirical test of difficulty detectability. On the other hand, the recency effect appeared to be slightly weaker for Experiment 1, suggesting another factor besides performance monitoring may have influenced sensorimotor confidence, although our efforts to quantify this effect were hampered by the nature of the error signals (see Supplementary information). An ideal test for use of a mixed strategy would involve keeping performance constant by fixing the difficulty while also varying likely uncertainty cues (e.g., titrating the mean and variability of the sensory signal; De Gardelle and Mamassian, 2015; Spence et al., 2015). This is more difficult in sensorimotor tasks as motor variability will introduce noise into the error signal, hindering any attempt to match performance. One way around this problem would be have participants judge sensorimotor confidence for replays of previously completed tracking and artificially adjust uncertainty cues. However, this would rely on metacognition acting similarly for active tracking and passive viewing, which has only been confirmed for motor-awareness confidence (Charles et al., 2020).

### 4.2 The recency effect

In the sensorimotor feedback process, incoming error signals inform upcoming action plans and quickly become irrelevant (Todorov, 2004; Bonnen et al., 2015). In contrast, the goal of performance monitoring for sensorimotor confidence is to accumulate error signals across time, much like the accumulation of sensory evidence for perceptual decisions with a fixed viewing time. In fact, in the accumulation-of-evidence framework, considerable effort has been made to incorporate a recency bias termed “leaky accumulation” (Busemeyer and Townsend, 1993; Usher and McClelland, 2001; Brunton et al., 2013; Matsumori et al., 2018). The main arguments for including a temporal-decay component is to account for memory limitations of the observer (e.g., from neural limits of recurrent excitation) or intentional forgetting for adaptation in volatile environments (Usher and McClelland, 2001; Nassar et al., 2010; Norton et al., 2019). For our task, memory constraints are a more likely explanation of the recency effect than intentional forgetting, because we have long trials of 6-14 s with no changes of stimulus statistics during a trial. One contributor to the error signal we have no control over, however, is the participant’s motivation to do the task. Even though tracking performance was constant when averaged across trials, fluctuations in motivation during a trial could lead to fluctuations in motor performance that do cause volatility in the error signal. Thus, alternating between bouts of good and poor performance could bias the participant to be more forgetful.

Previous efforts to characterise the time course of a metacognitive judgement have been limited to the perceptual domain. Using the reverse-correlation technique, Zylberberg et al. (2012) measured the temporal weighting function for confidence in two perceptual tasks and found a primacy effect: the initial hundreds of milliseconds of stimulus presentation had the greatest influence on perceptual confidence. Their finding and associated modelling suggests evidence accumulation for the metacognitive judgement stops once an internal bound for decision commitment has been reached. Our results suggest that sensorimotor confidence does not follow the same accumulation-to-bound structure, otherwise early error would have been more predictive of confidence than late error. One reason we may not have found a primacy effect is that the participant interacts with the stimulus to produce the errors that determine performance, allowing them a sense of agency that they can change or modify performance. As a result, there is no reason to settle on a confidence judgement based on initial performance. A contradictory finding to Zylberberg et al. (2012) is that sensory evidence late in the trial, during the period between the sensory decision and the metacognitive decision, can influence perceptual confidence in what is termed post-accumulation of evidence (Pleskac and Busemeyer, 2010), but this finding is hard to apply to our visuomotor task. Evaluating tracking is different from a single perceptual decision, because tracking is a series of motor-planning decisions (Wolpert and Landy, 2012). The error signal used to plan the next tracking movement is also the feedback of the error from the last moment of tracking. Additionally, subsequent estimates of target location could theoretically provide additional information about previous locations of the target. Identifying the source of the error signal for sensorimotor confidence, either by computational modelling or brain imaging, would help clarify the nature of the accumulation process.

So far we have considered an online computation of metacognition in parallel with sensorimotor decision making. Another alternative is that the evaluation of performance is computed retrospectively. Baranski and Petrusic (1998) showed that reaction times for confidence responses differed for speeded and unspeeded perceptual decisions, leading to the conclusion that perceptual confidence is computed online unless time pressure forces it to be evaluated retrospectively. It is reasonable to assume that the continual demand of cursor adjustment to track an unpredictable stimulus is taxing, leaving participants no choice but to introspect on their performance upon termination of the trial. If this were the case, we would likely see temporal biases consistent with memory retrieval. In the memory literature, there has been extensive evidence of both primacy and recency effects, which are thought to be associated with long-term and short-term memory processes respectively (Atkinson and Shiffrin, 1968; Innocenti et al., 2013). Thus, the observed recency effect in our experiment could be the result of short-term memory limitations constraining the time constant. Another reason observers may delay performance evaluation until after the trial is because tracking is typically a goal-directed behaviour, which can be evaluated by its success (e.g., catching the prey after a chase, hitting the target in a first-person shooter game, or correctly intercepting a hand in a handshake). Still, one may want to introspect about performance while tracking to decide whether the tracking was in vain. We did not incentivise participants to adopt a particular strategy in the task, so they may have treated error towards the end of the trial as their success in “catching” the target.

### 4.3 Metacognitive efficiency

We quantified metacognitive sensitivity for sensorimotor tracking with an AUROC metric that reflected the separation of the objective-performance distributions conditioned on sensorimotor confidence. This approach superficially shares some similarities with the metacognitive metric meta-*d*′ in perceptual confidence. For meta-*d*′, an ROC curve, relating the probability of a confidence rating conditioned on whether the observer was correct vs. incorrect, is computed as part of the analysis to obtain a bias-free sensitivity metric that reflects the observer’s ability to distinguish between correct and incorrect perceptual responses (Fleming and Lau, 2014; Mamassian, 2016). However, the area under this ROC curve (AUROC) has little meaning, as it is highly dependent on the sensitivity of the primary perceptual judgement (Galvin et al., 2003). Instead, the appropriate comparison is between the perceptual sensitivity, *d*′, and the metacognitive sensitivity, meta-*d*′. Typically, a ratio of these sensitivities is computed, with a value of 1 being considered ideal metacognitive efficiency (i.e., the best the observer can do given the identical sensory evidence available for the metacognitive judgement as the perceptual judgement). Empirically, ratios less than 1 are most often observed, indicating less efficient, more noisy decision-making at the metacognitive level (Maniscalco and Lau, 2012, 2016).

In contrast, the purpose of our AUROC metric is not to quantify how well the sensory information is used for the sensorimotor control versus sensorimotor confidence. Instead we used it as a non-parametric way of quantifying how sensitive an observer is to their true performance. The metric ranges from no sensitivity (i.e., chance performance) at 0.5 to perfect classification performance at 1. As with perceptual confidence, we do expect that the AUROC will depend to some degree on the variance in the performance of the primary task (e.g., tracking). For example, if there is little variance, then it should be difficult to identify well executed from poorly executed trials, whereas a large variance means performance could be more easily categorised. A second use of the AUROC metric was to quantify the degree to which a model of metacognitive behaviour could predict sensorimotor confidence (see Supplementary Information). By replacing the objective-performance axis with an internal decision-variable axis according to a model, a model’s explanatory power can be measured on a scale from none at 0.5 to perfect at 1. While we were unsuccessful at improving performance more than 5% in any of our experiments, which we did by accounting for both the recency effect and the effect of external sensory noise instead of simply computing RMSE using the true target location, the method of analysis nicely complemented our goal of quantifying how well sensorimotor confidence reflected objective performance.

We instead examined metacognitive efficiency by determining what error information contributed to sensorimotor confidence. The recency effect we observed constitutes an inefficiency in that not all information used for the primary sensorimotor decision-making was used for the metacognitive judgement as was instructed. Based on the similarity in shape of the recency effect for the duration conditions of Experiment 2, we can conclude that efficiency is inversely proportional to the duration of tracking. However, given long, multi-action sequences, it is not that surprising to find that some part of the perceptual information about error is lost. Some amount of forgetting is likely advantageous in real-world scenarios. For future metacognitive studies of action, it would be informative to examine estimates of sensorimotor confidence during action and how sensorimotor confidence interacts with goal planning, explicit learning, and expertise. For example, it would be worthwhile to investigate how sensorimotor confidence relates to cognitive control functions such as switching or abandoning motor tasks (Alexander and Brown, 2010), or how athletes and novices judge sensorimotor confidence (MacIntyre et al., 2014).

### 4.4 Conclusion

In sum, we found considerable evidence that humans are able to compute sensorimotor confidence, that is, they are able to monitor their motor performance in relationship to a goal. However, they do so inefficiently, in particular because of the recency effect that we revealed, disproportionately weighting the tracking error at the end of the trial to judge whether their performance was better than average. We replicated this effect with unpredictable stimulus-presentation durations to confirm that it was not the result of a response-preparation strategy. In our analyses, we have introduced the AUROC statistic, which we found useful for two purposes. First, it allowed us to quantify the relationship between sensorimotor confidence and objective tracking performance, and second, it provided a model-fit metric for elaborated decision models. Our results, obtained from a relatively simple goal of visuo-motor tracking, raise many questions for future studies on sensorimotor confidence. For example, is the recency effect a general phenomenon for sensorimotor confidence? And, does it result from online evidence accumulation of error or a retrospective memory retrieval upon termination of movement? What factors determine the strength of the recency effect for sensorimotor confidence (i.e., attention, sensorimotor goals, etc.)? Further work will help provide a clearer link between models of sensorimotor behaviour and models of motor metacognition.

## Acknowledgements

We thank Eero Simoncelli for fruitful discussions that contributed to this project. This work was supported by NIH Grant EY08266 and National Science Foundation Collaborative Research in Computational Neuroscience Grant 1420262, as well as French ANR grant ANR-18-CE28-0015-01”VICONTE”.

## Author note

This research was presented at the 2017 Vision Sciences Society meeting at St. Pete Beach, FL and the 2017 Conference on Cognitive Computational Neuroscience in New York, NY. Data from this study can be found at https://osf.io/enxdt/. Individual author contributions in the CRediT taxonomy style are as follows (black: major contribution, gray: minor contribution, see https://www.casrai.org/credit.html for more details):

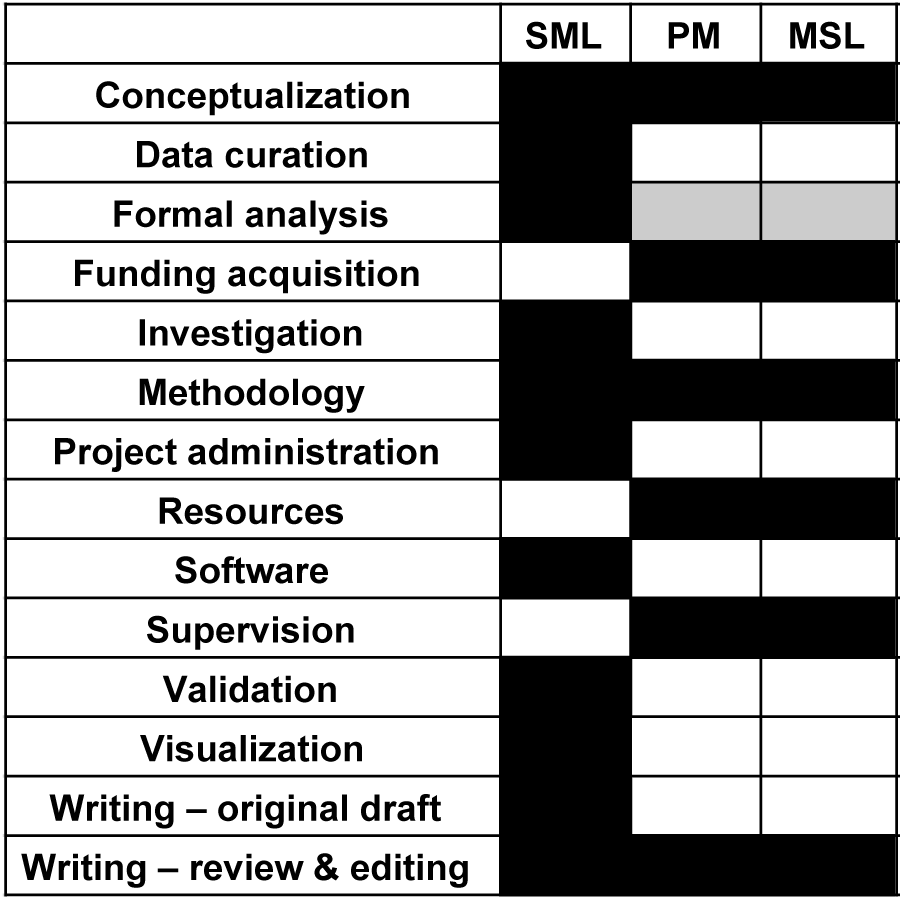

