## Supplementary Information for "Performance Monitoring for Sensorimotor Confidence: A Visuomotor Tracking Study"

### 1 Models of sensorimotor confidence

The goal of the modelling was to quantify the recency effect for sensorimotor confidence observed in both experiments to facilitate comparison. First we established whether a model of the recency effect would better fit the data than other plausible candidate models of sensorimotor-confidence behaviour. We considered a total of four models of how an observer might extract a decision variable from this error signal for sensorimotor confidence. The *RMSE model* uses the RMSE statistic calculated from the tracking error. In the *recency model*, absolute tracking error was exponentially weighted, i.e., the decision variable was $x = \int_0^{t_{\text{final}}} |err(t)| e^{-(t_{\text{final}}-t)/\tau} dt$, where the time constant $\tau$ was an additional free parameter. In the other two models, *max-error* and *final error*, we considered the sub-optimal strategies of simply taking the maximum absolute error or the final error (equivalent to $\tau \to 0$) as the decision variable. For all models, we used a cumulative log-normal linking function with two free parameters (bias and slope; lapse rate was fixed at 2%) to convert the continuous internal estimate of error into a binary sensorimotor-confidence judgement of "better" or "worse". This linking function was selected because the decision variable could not be less than 0. We performed the analysis twice using different tracking-error time series each time: the horizontal error between 1) the cursor and target, and 2) the cursor and perceptual estimate as per the smoothing model of Figure 6A.

All models were fit using custom MATLAB scripts that calculated the maximum likelihood estimates (MLEs) of the parameters using gradient decent. The log-likelihood values at the MLE were then converted to corrected Akaike information criterion (AICc) scores for model comparison (Akaike, 1974; Cavanaugh, 1997), which penalises the number of parameters in each model. The AICc was used instead of the uncorrected form because this metric is better suited to cases with small sample sizes (Cavanaugh, 1997). We also used a second method of model comparison: the AUROC statistic computed using the internal estimate of error according to each model, which reflects how well a model predicts the sensorimotor confidence. The AUROC would be 1 for a model that perfectly captures sensorimotor confidence, whereas a model that does not will have an AUROC closer to 0.5.

#### 1.1 Experiment 1

The results of the AICc model comparison for Experiment 1 are shown in Figure S1A, with all models compared to the recency model. For both the cloud-size and the velocity-stability session, the recency model is best supported by the data. However, the RMSE model also fit quite well. The relative AICc scores for these two models with the input of objective error were $\Delta\text{AIC}_c = 6.35 \pm 4.33$ and $\Delta\text{AIC}_c = 4.85 \pm 4.10$, for cloud-size and velocity-stability sessions respectively. Similar results were obtained when using perceptual error for the RMSE model, as the relative scores were $\Delta\text{AIC}_c = 4.34 \pm 2.83$ and $\Delta\text{AIC}_c = 3.41 \pm 3.41$. The maximum-error model fit less well, and the final-error model error fit behaviour very poorly. We also refit the models to the data omitting the first 2 s of tracking to rule out any abnormal weighting at the start the trial and the pattern of results was unchanged. Additionally, how well each model predicted sensorimotor confidence was quantified by calculated the AUROC using the model's internal decision variable (Figure S1B). For both objective and perceptual error, there was a marginal improvement for the recency models over the maximum-error model in both sessions and over the RMSE model for the cloud-size session but not the velocity-stability session (at

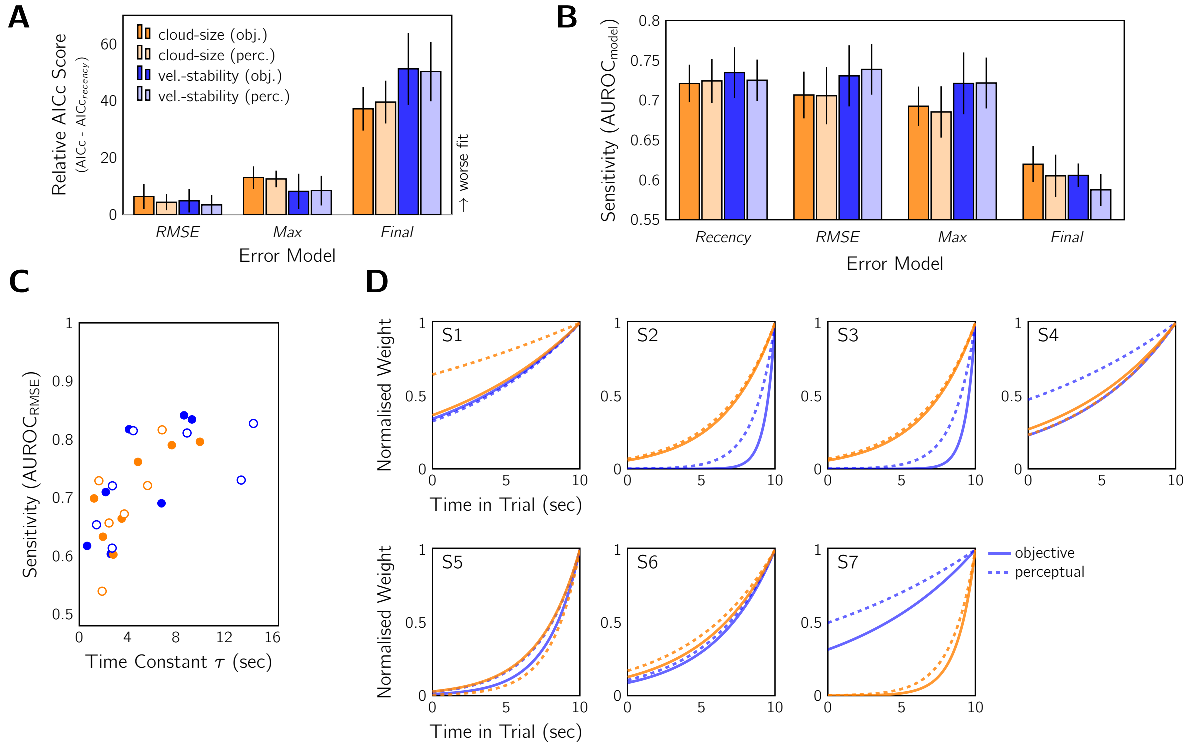


Figure S1: Decision models for sensorimotor confidence in Experiment 1 fit considering objective error and perceptual error (see text for details). The four models were 1) recency: absolute tracking error was weighted according to an exponential function to give a recency-like effect; 2) RMSE: correctly estimates the tracking RMSE; 3) maximum error: only the largest error determined confidence; and 4) final error: only the error at the end of tracking was considered. A: Model-comparison results using the AICc scores relative to the recency model (mean $\pm$ SEM across participants), which penalised model complexity. All other models were worse predictors of the subjective evaluations than the recency model. B: Same as in (A), but showing AUROC according to each model type. Higher AUROCs indicate that model is a better predictor of sensorimotor confidence. C: Relationship between the time constant $\tau$ from the recency model and the metacognitive sensitivity AUROC as per the RMSE model (reported in the main paper). Datapoints: individual participants and sessions (indicated by colour). Fill: error type either objective (filled) or perceptual (unfilled). D: Best-fitting temporal weighting functions for the recency model for each subject, session (colour), and error type (line type). Weighting functions were normalised to have a final weight of 1.

most $\sim 5\%$). The final-error model was a poor predictor of sensorimotor confidence. In general, this pattern of results aligned with the AICc scores in Figure S1A.

Figure S1D shows the best-fitting exponential weighting function for each observer in each condition. There is a high degree of consistency between the time constants of the sessions for all but two observers, which also differed from each other in terms of which session had the longer time constant. On average, $\tau = 4.6 \pm 1.2$ s (mean±SEM) for the cloud-size condition and $\tau = 4.9 \pm 1.3$ s for objective error, and $\tau = 6.4 \pm 2.8$ s and $\tau = 6.9 \pm 2.0$ s for perceptual error. Figure S1C shows the relationship between the best-fitting $\tau$ and AUROC reported in the main document (i.e., AUROC for the RMSE model with objective error) for each of the sessions.

To quantify the relationship between the recency model and the metacognitive sensitivity results, we performed a linear mixed-effects (LMM) analysis. The dependent variable was the AUROC statistic from the RMSE model. The fixed effects were the time constant $\tau$, session (cloud-size, velocity-stability), error type (objective, perceptual), and an intercept term. Participant was the random effect. We found a significant positive relationship between $\tau$ and the AUROC statistic, according to a Type II Wald chi-square test ($\chi^2 = 6.31$, $p < .05$). A positive relationship is expected because the larger $\tau$ is, the closer the temporal weighting is to the calculation of the RMSE used in the ROC-style analysis. There was no significant effect of session or error type, nor were there any two-way or three-way interaction effects (all $p > .05$). Taking just the objective-error results, when the recency model was fit with a single time constant for both difficulty manipulations instead of a separate time constant for cloud-size and velocity-stability sessions, the model fit was only marginally impaired (AICc$= 584.79 \pm 15.57$ for combined versus AICc$= 577.02 \pm 14.46$ for separate) and $\tau = 4.52 \pm 1.03$.

#### 1.2 Experiment 2

Our analysis of Experiment 2 repeated what was done for Experiment 1, but instead had two recency models: 1) a single time constant for all duration conditions, or 2) a

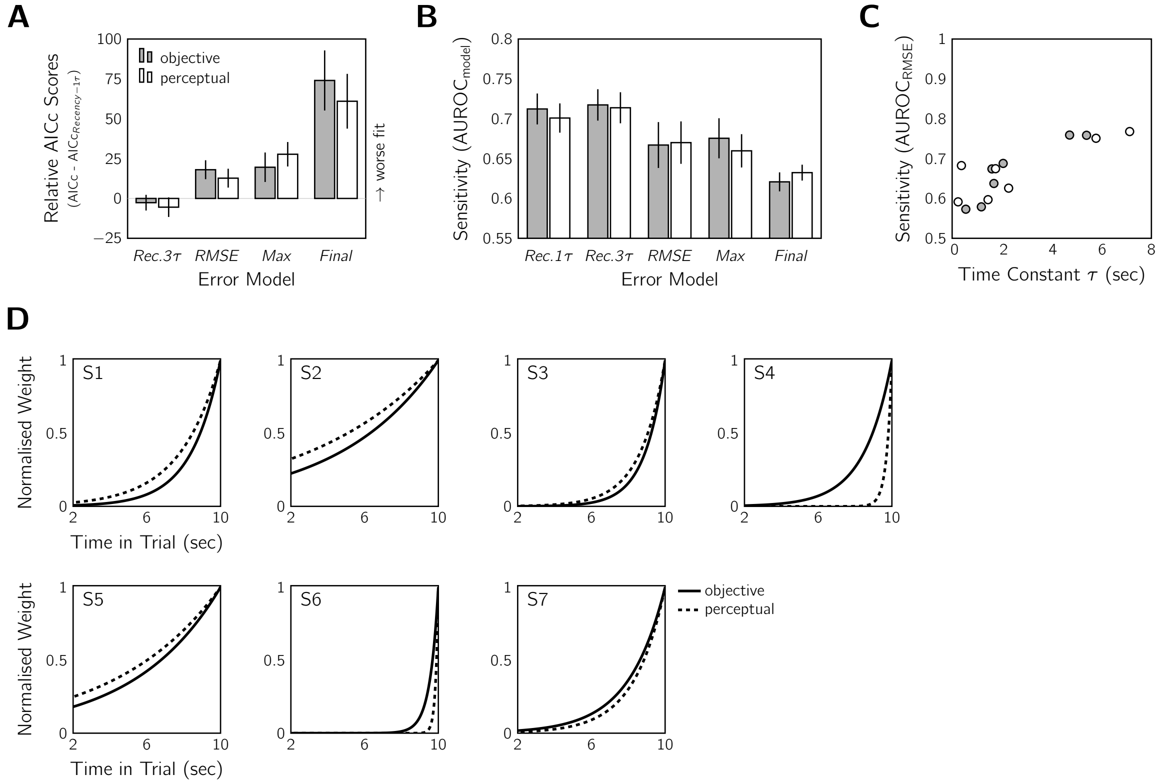


Figure S2: Decision models for sensorimotor confidence in Experiment 2 fit considering objective error and perceptual error (see text for details). We examined the same four models as for Experiment 1 with an additional fifth model, recency-3$\tau$, which was the recency model fit with an independent $\tau$ parameter for each duration condition. A: Model-comparison results using the AICc scores relative to the recency model with a single $\tau$ (mean $\pm$ SEM across participants). Only error after 2 s was considered. The recency models were fairly comparable and all other models performed worse. There was little difference if we used objective (grey) or perceptual (white) error. B: Same as in (A), but showing AUROC according to each model type. Higher AUROCs indicate that model is a better predictor of sensorimotor confidence. C: Relationship between the time constant $\tau$ from the two recency models and the corresponding metacognitive sensitivity AUROC from the RMSE model. Data points: individual participants. Error type: objective (grey) or perceptual (white). Note the change in x-axis scale from Figure S1C. D: Best-fitting temporal weighting functions for the single-$\tau$ recency model with results from both error types shown (objective: solid; perceptual: dashed). Weighting functions were normalised to have a final weight of 1. Note the change in x-axis scale from Figure S1D as error before 2 s was ignored for all model fits.

separate time constant for each duration condition. The three other models, which had no additional free parameters, were the same as for Experiment 1. Model fits were also assessed with the same model-comparison methods. As shown in Figure S2A, the best-fitting models were the recency models, with the $3\tau$ model fitting slightly better than the single-$\tau$ model (objective error: $\Delta\text{AIC}_c = -2.65 \pm 4.92$; perceptual error: $\Delta\text{AIC}_c = -5.47 \pm 6.14$). Again, we calculated how well the model predicted sensorimotor confidence by finding the AUROC according to each model type (Figure S2B). There was a marginal improvement for the recency models over the RMSE model. The maximum-error model gave comparable results to the RMSE, and the final-error model was the worst predictor. In general, this pattern of results aligned with the AICc scores in Figure S2A. Figure S2C shows the $\tau$ parameter estimates from the single-$\tau$ model contrasted against metacognitive sensitivity for each participant. We found a significant positive relationship between $\tau$ and metacognitive sensitivity with a LMM analysis (fixed effects: $\tau$, error type, intercept; random effect: participant) according to a Type II Wald chi-square test ($\chi^2 = 9.37$, $p < .01$). There was no significant effect of error type, but there was a significant interaction between error type and $\tau$ ($\chi^2 = 6.82$, $p < .01$). On average, $\tau = 2.38 \pm 0.71$ s for fitting with objective error and $\tau = 2.64 \pm 1.03$ s with perceptual error, both of which are about half a long as the time constants observed in Experiment 1. The temporal weighting functions of individual participants are shown in Figure S2D. Notice that the perceptual and objective error tend to yield a similar time constant.

#### 1.3 Limitations of the analysis

Finally, we wish to highlight some limitations of this modelling approach and caution against overinterpreting the specific values of $\tau$ obtained. Despite the reasonable decision-function fits (Figure S3A) and high within-subject consistency for the $\tau$ parameter for Experiment 1, it was difficult to calculate confidence intervals for the parameter estimates for both experiments. Error bars were very wide when model fits were analysed with non-parametric bootstrapping or even brute-force calculation of the marginal likelihood.

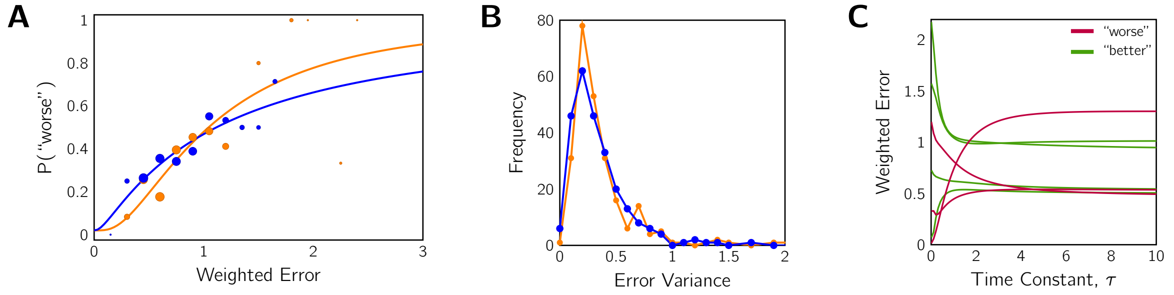


Figure S3: Limitations of the modelling approach illustrated using an example participant from Experiment 1. A: Example best-fitting decision-linking functions. Data points: binned proportion of trials participant responded "worse", with marker size scaled by the log of the number of trials. Solid lines: model fit with MLE estimates. B: Distribution of the within-trial tracking-error variance for each condition. C: Seven example trials illustrating how weighted error varies as a function of the smoothing parameter $\tau$, coloured by metacognitive response. There is very little weighted-error variability for time constants more than 4 s when considering individual trials.

For example, in the bootstrapping analysis of Experiment 1, the distribution of best-fitting $\tau$ across sampled datasets was multimodal, with up to a quarter of datasets being fit best by a much larger $\tau$ or hitting the reasonable bounds of the parameter space. Figure S3B shows a potential reason for this sample-driven problem: within-trial error variance differences between trials. That is, trials with higher variability will contribute more to constraining the estimate of $\tau$ than those with lower variability. Which, if any, of these higher-variance trials are sampled for the bootstrapping can disproportionately affect the modelling outcome. An additional issue, affecting even the brute-force attempt to get confidence intervals, is the way $\tau$ and weighted error interact (see Figure S3C for some examples). Time constants larger than about 4 s all produce approximately the same weighted error and consequently will all fit similarly well. Thus, distinguishing the error-weighting model with certainty is not feasible unless the participant is using a shorter time constant. This was not the case with more than half of our participants.

**Summary:** The research goals that prompted us to fit decision models for the sensorimotor confidence were to 1) determine whether a recency-effect model did indeed best explain the confidence judgements, and 2) quantitatively compare the recency effects in Experiments 1 and 2. For the first goal, a simple model comparison showed that the recency-effect model, with an exponential weighting function for tracking error, was superior to several other candidate models in explaining the results of both experiments.

However, all models except the final-error model fit similarly well for Experiment 1. In contrast, the recency models of Experiment 2 were clearly superior to all other models, both in the AICc comparison and the AUROC comparison, with about a 5% gain in predictability from consideration of the recency effect. Our results changed very little if we considered objective tracking error or perceptual error that accounted for external noise. As for the second goal, we found that the time constants that reflected the strength of the recency effect differed between experiments, with a stronger effect (i.e., smaller $\tau$) for the fixed-difficulty, mixed-stimulus-duration task of Experiment 2. Though we caution against overinterpreting the parameter estimates. While our analyses might appear as a decisive win for the recency model, there are limitations in the datasets preventing a clear quantitative answer. For this reason, we chose to present these analyses here rather than in the main paper.
